# Embodied Emergence of Temporal Complexity in Finger Tapping: Synaptic and Dopaminergic Control in Cortico-Basal Ganglia Loops

**DOI:** 10.64898/2026.09.07.749738

**Authors:** Andrea Tigrini, Haruki Shimokado, Kazuki Matsui, Alessandro Mengarelli, Rami Mobarak, Mara Scattolini, Federica Verdini, Laura Burattini, Taishin Nomura

**Affiliations:** Università Politecnica delle Marche, Department of Information Engineering, Via Brecce Bianche, 12, 60131, Ancona; Kyoto University, Graduate School of Informatics, 606-8501 Yoshida Honmachi, Sakyo-ku, Kyoto; Università eCampus, Department of Theoretical and Applied Science, 22060, Novedrate, Italy

## Abstract

Finger tapping is a crucial clinical window into the brain’s motor integrity, particularly in Parkinson’s disease (PD). Beyond mere rhythmicity, healthy motor output exhibits long-range correlations (LRCs)—a hallmark of temporal complexity and physiological adaptability. However, how this fractal structure emerges from the interplay between neural circuits and physical body dynamics, and why it collapses under dopamine depletion, remains elusive. Here, we present an embodied computational model integrating the cortico-basal ganglia-thalamocortical (CBGT) loop with a physical finger model under intermittent control. We demonstrate that LRC is not stochastic noise, but emerges from the interaction between physical finger dynamics and intermittent, sensory-driven action selection. Our simulations reveal that while unbiased action selection yields random variability, a sensory-driven striatal bias generates robust LRC, demonstrating that motor complexity can be actively driven by neural “synaptic bias” rather than passive body mechanics. Crucially, simulating PD pathology via dopamine reduction disrupts this bias, leading to a loss of complexity and white-noise transitions that recapitulate a key qualitative feature of the fractal breakdown observed in PD. By bridging neuro-modulation, biomechanical execution, and macroscopic behavior, this work provides a mechanistic framework for embodied neural information processing and offers a potential digital-twin platform for objective biomonitoring in neurodegenerative disorders.

**Author summary:** Finger tapping may look simple, but the timing of each tap carries rich information about how the brain and body work together. In healthy movement, small variations from one tap to the next are not completely random; instead, they show long-range patterns that are often reduced in Parkinson’s disease. We asked how these patterns arise and why they may disappear when dopamine levels fall. To address this question, we built a computer model that combines neural circuits involved in movement selection with the physical mechanics of the finger. We found that realistic temporal patterns emerged only when sensory information from the moving finger biased the brain’s selection of the next action. When this bias was removed, the tapping rhythm became much more random, even though tapping itself continued. We also found that reducing dopamine progressively disrupted these long-range patterns, reproducing an important qualitative feature of motor impairment in Parkinson’s disease. Our results suggest that healthy movement complexity emerges from continuous interaction between neural decision-making and body mechanics. This framework may eventually help link simple movement measurements to changes in underlying neural function.

## 1 Introduction

Repetitive motor tasks, such as finger tapping and gait, are governed by complex neural architectures that ensure both stability and flexibility [1]. A defining characteristic of healthy motor systems is the presence of long-range correlations (LRC) in cycle-to-cycle variability [2]. This positive persistency—where the scaling exponent *α* ≈ 1 in Detrended Fluctuation Analysis (DFA)—signifies a temporal “memory” spanning hundreds of cycles, reflecting a state of optimal adaptability [3]. LRCs were studied extensively in gait cycle variability [4, 5] as well as in inter-tap time interval during finger tapping [6–10] with mathematical timeseries modeling [8, 11]. In neurodegenerative disorders such as Parkinson’s disease (PD), this temporal complexity is profoundly disrupted [4, 12]. PD patients, particularly those prone to freezing of gait [4] or freezing of tapping [13], exhibit a shift toward more random, white-noise variability (*α* ≈ 0.5) [4, 14]. This loss of complexity is more than a passive symptom; it is a direct manifestation of an unstable, rigid motor control system unable to maintain flexible rhythmic output [2]. Mechanistically elucidating the etiology of this fractal breakdown is critical for developing non-invasive dynamical biomarkers and *in silico* platforms for early warning and disease progression [2, 15, 16].

Current neurological consensus points to the cortico-basal ganglia-thalamocortical (CBGT) loop as the primary engine for action selections during finger tapping rhythmogenesis [17–20]. The continuous competition between the direct (Go), indirect (NoGo), and hyperdirect (Brake) pathways determines how specific motor programs, such as finger extension and flexion, are selected and executed [21–23]. While sophisticated spiking neural networks have attempted to model these neuronal dynamics, they frequently focus on static connectivity, missing the dynamic, automatic decision-making inherent to continuous rhythmic execution [24–26]. Recent advances, such as the Ursino model [20], have introduced computationally efficient “neural mass” descriptions of the CBGT loop, highlighting the intermittent nature of motor control where cortical motor commands are applied only when an evidence-accumulation threshold is reached.

However, a significant gap remains at the intersection of neuroscience and biomechanics: these models typically lack a representation of the physical effector [19, 20, 26, 27]. Since sensory feedback regarding the mechanical state of the body is a primary input to the striatum [28], the absence of a physical body limits our understanding of how neural competition for action selection is grounded in physical reality. Crucially, while previous posture-control studies suggested that temporal complexity can emerge from passive mechanical biases of the body [29], how fractal dynamics are actively sustained during volitional, intermittent movements remains unknown. Indeed we may wonder that, without this tight neuromechanical coupling, which ensures embodied neural information processing, stochastic neural models alone could fail to reproduce the robust LRC observed in humans.

Thus, in this study, we present an integrated *in silico* framework that couples a CBGT intermittent control model with a physical biomechanical model of the index finger for modeling the emergence of LRC in cycle-to-cycle variability during periodic finger tapping. We hypothesize that LRC is not generated by stochastic neural noise but emerges from the active accumulation of sensory-biased evidence within cortico-striatal pathways interacting with physical execution. By systematically manipulating dopamine levels and network weights, we demonstrate that the presence of LRC is highly sensitive to the degree of striatal selection bias.

Our findings reveal that the dopamine depletion characteristic of PD directly impairs this active mechanism, shifting the system from an adaptable fractal state to random white noise. By bridging synaptic-level neuromodulation, biomechanical execution, and macroscopic behavioral complexity, this work provides a mechanistic framework for embodied motor control and establishes a potential digital-twin platform for objective biomonitoring in neurodegenerative disorders.

## 2 Results

### 2.1 Experimental long-range correlations in human self-paced finger tapping

An example of self-paced finger tapping in a healthy young adult is shown in **Figure 1**. Here, the inter-tap interval (ITI)—defined as the time between two consecutive finger extension peaks—was analyzed as a behavioral motor time series over a 250-second trial (**Fig. 1 left**). The DFA scaling exponent *α >* 0.5 confirms that the ITI time series exhibits LRC with persistent behavior (**Fig. 1 right**), as reported by many previous studies [6–10]. This demonstrates that the oscillatory alternation of flexion and extension during self-paced tapping is not random noise; rather, like other cyclical human movements, it is orchestrated by complex, multi-scale neural activity.

**Fig 1.**
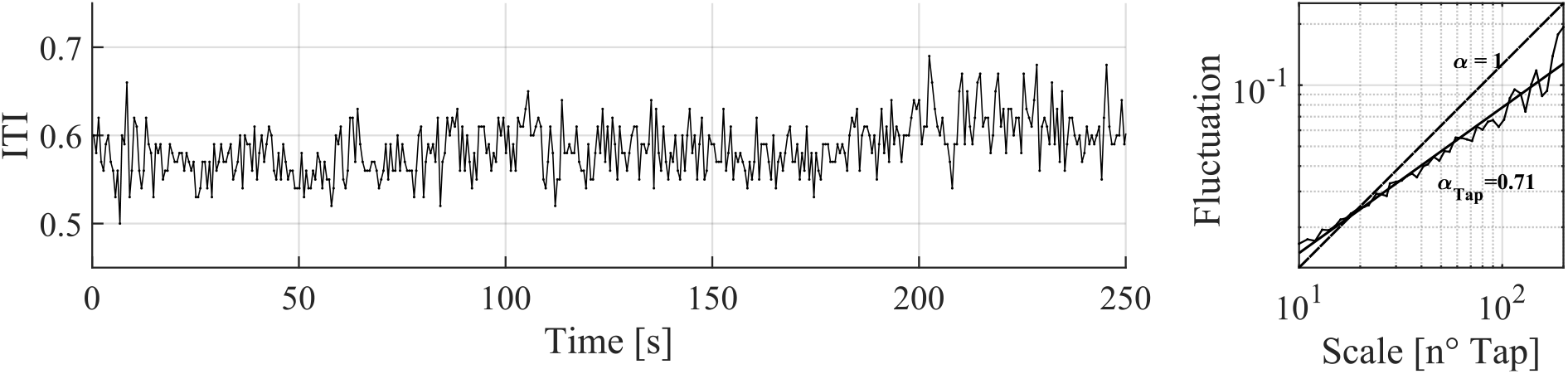
Long-range correlations in human self-paced finger tapping. Left panel: Representative time series of the ITI obtained from a healthy young adult during a 250-second self-paced finger tapping task. Right panel: DFA of the ITI time series. The log-log plot of fluctuation *F* (*n*) versus window size *n* reveals a linear relationship with a scaling exponent *α >* 0.5, indicating persistent long-range correlations.

To understand how this complex dynamic behavior arises from the interaction between central neural circuits and peripheral biomechanics, a biologically plausible model must be capable of reproducing these long-range correlation properties when its parameters are properly tuned.

### 2.2 A neuromechanical model of sustained periodic finger tapping

Despite the apparent simplicity and automaticity of finger tapping, accumulating evidence suggests that the CBGT loop plays a pivotal role in rhythmogenesis [17, 18]. Specifically, it mediates sequential decisions regarding when and which index finger muscle—the extensor (E) or the flexor (F)—the cortical motorneurons should activate to execute structured extension or flexion.

Figure 2 illustrates the overall architecture of the embodied model developed in this study, which builds upon the neuroanatomical structure of the CBGT network implemented in the neural mass formulations by Ursino et al. [20]. As in general motor control tasks, the fundamental role of the basal ganglia (BG) within the CBGT loop is to sequentially select the appropriate muscle for a required motor task. In this study, this corresponds to choosing between the antagonistic muscle pairs of the index finger. The loop subsequently determines whether to contract (Go) or suppress (NoGo) each muscle by modulating the direct (Go) or indirect (NoGo) pathways. To achieve sustained, spontaneous, periodic tapping, the motor cortex (MC) undergoes a continuous sequence of action-selection decisions utilizing two distinct cortical neural mass models (hereafter referred to as cortical neurons): the cortical E neuron driving extension, and the cortical flexor F neuron driving flexion.

**Fig 2.**
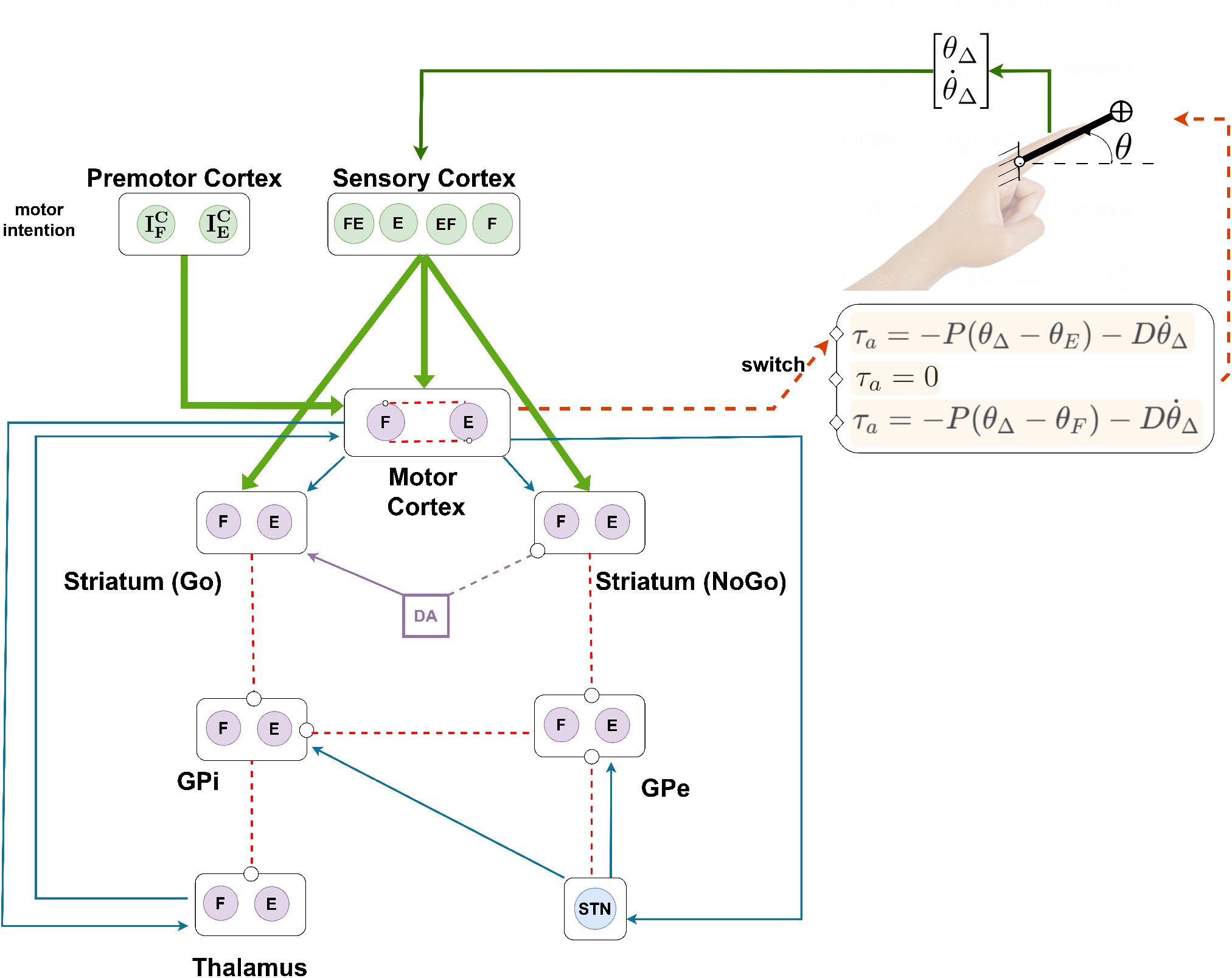
General scheme of the neuromechanical model used in the experiments. BG are represented as compartments, where each neuron is associated with a specific action, i.e., E or F. The STN is modeled as a single-neuron compartment. Cyan solid arrows indicate excitatory synapses, while red dashed lines ending with circles indicate inhibitory synapses. The violet DA box highlights the role of this parameter in the Go and NoGo pathways of the striatum. Sensory information flow and coding are shown with green arrows, while orange arrows indicate the intermittent control actions selected by the MC compartment.

Figure 3 (top trace) exemplifies the oscillatory, alternating activation of these two cortical neurons, following drift-diffusion dynamics [30]. The activation level of each neuron represents the dynamic accumulation of sensory and looped evidence toward decision-making. The evidence accumulated by the two cortical neurons competes continuously; the neuron that first reaches a predefined decision threshold wins the competition, selecting the corresponding action in a winner-takes-all manner. This WTA characteristic is enforced via reciprocal inhibitory connections between the two cortical populations (see Fig. 2). Once the cortical E neuron wins the competition and remains suprathreshold, an active torque *τ*_*a*_—modeled by a proportional-derivative feedback controller—exerts an active extensor torque (*τ*_*a*_ = *τ*_*E*_) on the finger joint to drive it toward the desired extension angle *θ*_*E*_. Conversely, when the cortical F neuron wins and remains suprathreshold, an active flexor torque (*τ*_*a*_ = *τ*_*F*_ ) drives the finger toward the desired flexion angle *θ*_*F*_ . The target configurations of *θ*_*E*_ and *θ*_*F*_ within the 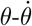 phase plane, which describes the mechanical state of the finger, are illustrated in Figure 4. Specifically, the active feedback control torques for extension and flexion are proportional to the position errors (*θ*_Δ_ − *θ*_*E*_) and (*θ*_Δ_ − *θ*_*F*_ ), respectively, combined with a derivative term proportional to 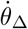. Here, *θ*_Δ_ and 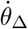 represent the joint angle and angular velocity incorporating a delay in sensory feedback of Δ seconds.

**Fig 3.**
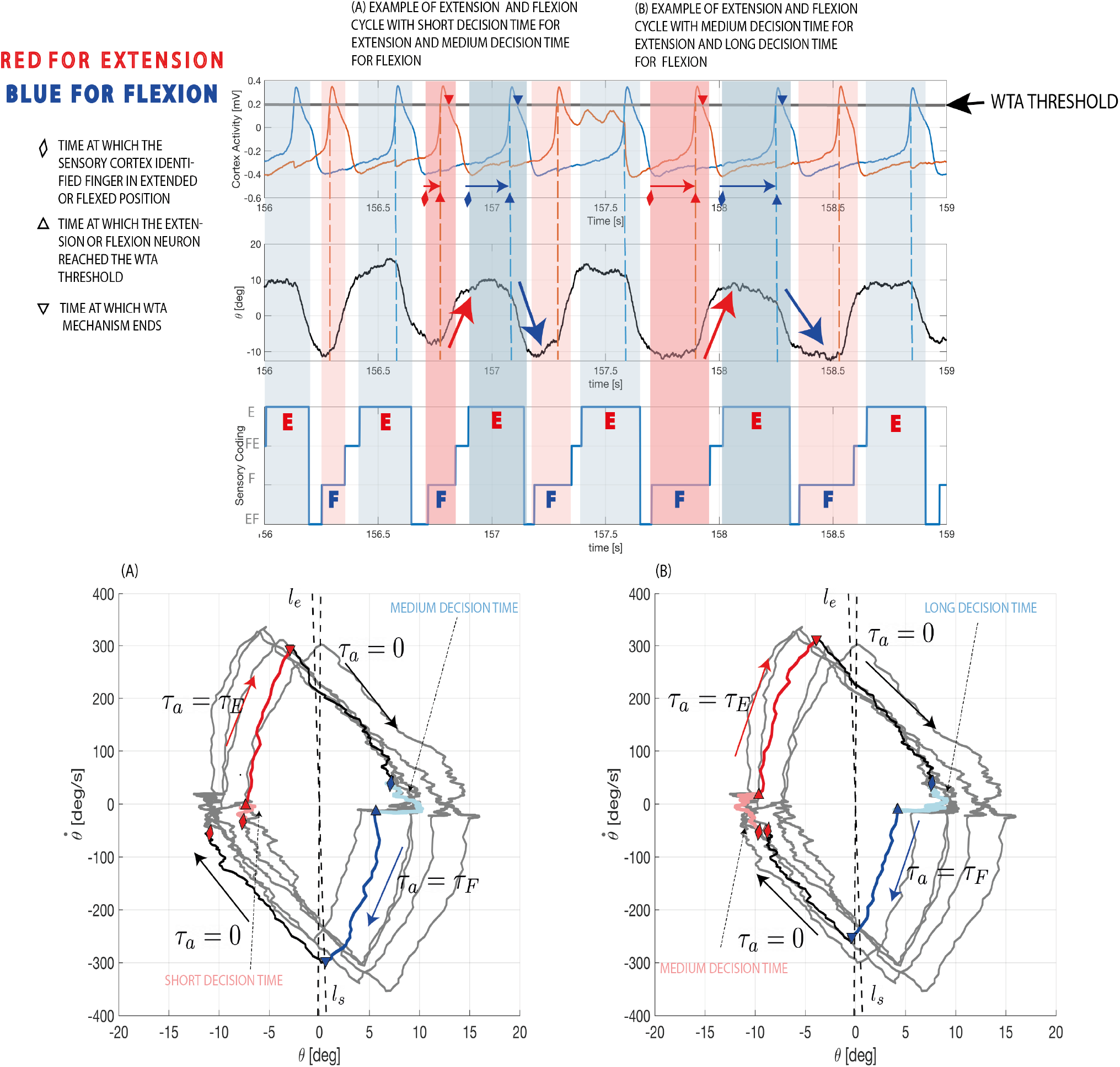
Time series of cortical activity, finger sway angle, and sensory coding states over 3 seconds of simulation with DA = 0.5. Two examples of extension E-F cycles are highlighted, and the corresponding finger dynamics are shown in the phase portraits in the bottom panels (A) and (B). In the top panel, the winner-takes-all (WTA) threshold is indicated by a solid black line in the MC activity. Red and blue diamonds mark when the sensory cortex encodes the finger position in E or F states, triggering neural competition. Up-pointing triangles indicate the time at which the E neuron (red) or F neuron (blue) wins the competition. Similarly, down-pointing triangles indicate the end of the WTA mechanism. The time from a diamond to the up-pointing triangle defines the decision time, which is highlighted in the phase portrait panels by light red and light blue segments of the trajectories. The time between up- and down-pointing triangles defines the interval during which the active torque *τ*_*a*_ = *τ*_*E*_ or *τ*_*a*_ = *τ*_*F*_ is applied, corresponding to the solid red and blue segments of the trajectories in panels A and B, respectively.

**Fig 4.**
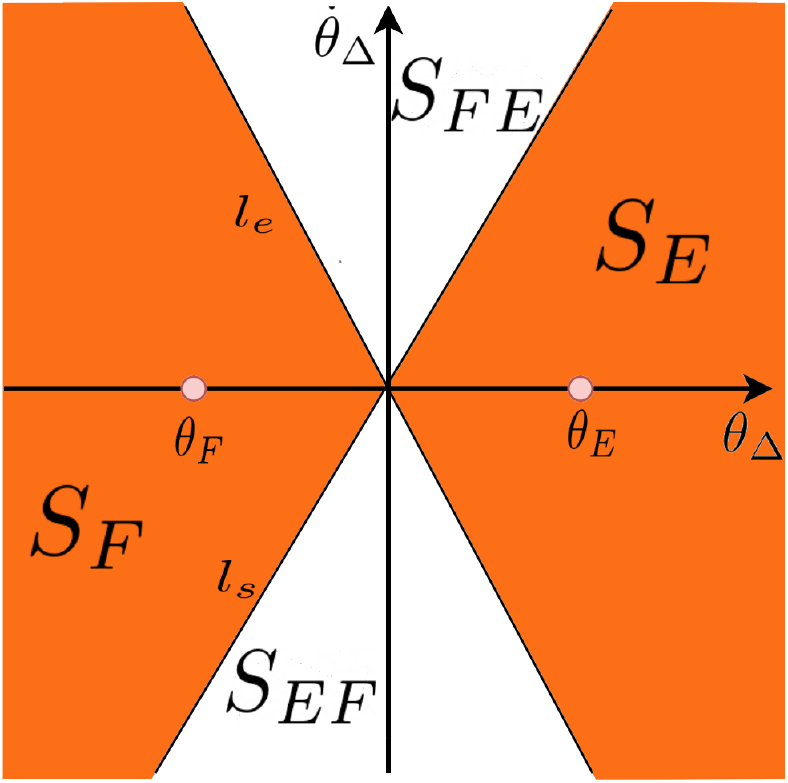
General partitioning scheme for sensory cortex coding, on the phase portrait of the delayed variable *θ*_Δ_ and 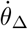. The regions are mathematically expressed as reported in eq. (6)

Crucially, no active control torque is applied to the finger during the subthreshold competition process until either the cortical E or F neuron crosses the threshold, defining the decision time (DT). Each DT serves as a “control-off” period for the feedback controller, during which the finger moves passively, driven solely by inertial forces and torque noise. This architecture inherently yields an intermittent control scheme. Motor behavior generated by this intermittent control is visible as the meandering trajectory of the finger state in the 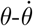 phase plane, associated with a null active control torque (*τ*_*a*_ = 0) near *θ* = *θ*_*E*_ and *θ* = *θ*_*F*_ (see Figure 3, lower panels A and B). These steady-state kinematics are generated because the active control torque drops to zero during each DT of the drift-diffusion-like mechanism—namely, when the activities of the cortical E and F neurons compete within their sub-threshold regimes.

Figure 3 (second trace) displays the simulated joint angle of the finger, illustrating alternating extension (upward) and flexion (downward). Critically, the timings at which the cortical E neuron activity reaches the decision threshold coincide closely—albeit stochastically—with the kinematic reversal points from flexion to extension (compare the top and second traces of Figure 3). Similarly, the threshold-crossing timings of the cortical F neuron match the reversals from extension to flexion. A prolonged DT for the extensor or the subsequent flexor manifests as a lengthened ITI, whereas abbreviated DTs yield shorter ITIs. Consequently, LRC properties of the system are predominantly dictated by the macroscopic sequence of these micro-scale DTs.

We modeled the cortical neurons using a Class 1 Morris-Lecar formulation [31, 32]. The synaptic input to each cortical neuron comprises CBGT-mediated thalamic signals, sensoricortical feedback signals, and tonic current injections from the premotor cortex. As noted above, the rise in membrane potential represents the accumulation of decision evidence. The cortical neuron models are tuned such that the total synaptic input fluctuates near a saddle-node-on-invariant-circle (SNIC) bifurcation point [31]. Consequently, modulations in synaptic input are faithfully translated into changes in firing frequency (governed by a square-root relationship with the synaptic input) [31], which directly modulate the inter-decision intervals and the resulting ITIs. Under this formulation, if the variability of the synaptic input to the cortical neurons exhibits LRC, the ITIs reflect this correlation accordingly. When the synaptic input exceeds the bifurcation threshold, the dynamics of the cortical neurons become intrinsically oscillatory [31], rendering the selection of finger extension and flexion fully automatic and driven by coupled limit-cycle oscillators. Therefore, operating near the bifurcation point allows the system to perform decision-making at the critical boundary between intentional choice and automatic response.

The primary driver of the synaptic input to each cortical neuron is the excitatory thalamo-cortical signal, which is gated by inhibitory projections from the globus pallidus internus (GPi) to the thalamus; hence, thalamic output is enhanced when GPi activity is suppressed (disinhibition). This disinhibition is regulated by the magnitude of excitatory inputs from the sensory cortex to the striatum. In our model, the mechanical state of the finger is fed back into the CBGT circuit via its representation in the sensory cortex, where the instantaneous state of the finger is categorically encoded by four distinct sensory cortical neurons designated as FE, E, EF, and F (see the sensory cortex layer in Figure 2). Specifically, the finger’s 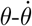 phase plane is partitioned into four functional regions, as illustrated in Figure 4. When the delay-affected state of the finger lies within the region *S*_*E*_, the sensory cortical E neuron is activated, emitting a binary signal of “1” to the striatal and cortical neurons, while the remaining sensory neurons remain quiescent (“0”). The corresponding state vector of the sensory cortex is denoted as *S* = [*S*_*FE*_, *S*_*E*_, *S*_*EF*_, *S*_*F*_ ] = [0 1 0 0]. Similarly, the FE, F, and EF sensory neurons are uniquely activated when the state point enters the *S*_*FE*_, *S*_*F*_, and *S*_*EF*_ regions, respectively. As the state point of the finger rotates clockwise in the phase plane during periodic tapping (Figure 3, lower panels), starting from an extended position, the sensory cortex exhibits cyclic state transitions: [0 1 0 0] → [0 0 1 0] → [0 0 0 1] → [1 0 0 0] (see the third trace of the upper panels in Figure 3).

### 2.3 Key mechanisms of action selection

When finger extension within the region *S*_*E*_ elicits the sensory state *S* = [0 1 0 0], the cortical sensory E neuron delivers excitatory drives to the striatal direct pathway neurons for both the flexor (Go-F) and the extensor (Go-E). The magnitudes of these inputs are governed by the corticostriatal synaptic weights projecting from the sensory E neuron to the Go-F and Go-E neurons, denoted as *W*_*GS*_ in Figure 5(A)-1. Heightened activity in the Go-F (or Go-E) neuron inhibits the corresponding GPi-F (or GPi-E) population, thereby disinhibiting the thalamic F (or E) neuron. The resulting surge in thalamic drive to the cortex facilitates the onset of finger flexion (or extension). Concurrently, the selection of antagonistic actions is regulated by synaptic projections from the sensory E neuron to the striatal NoGo neurons (*W*_*NS*_ in Figure 5(A)-1). Identical computational principles govern the loops initiated by sensory drives to the Go and NoGo populations from the sensory neurons representing states *S*_*EF*_ (Figure 5(A)-2), *S*_*F*_ (Figure 5(A)-3), and *S*_*FE*_ (Figure 5(A)-4).

**Fig 5.**
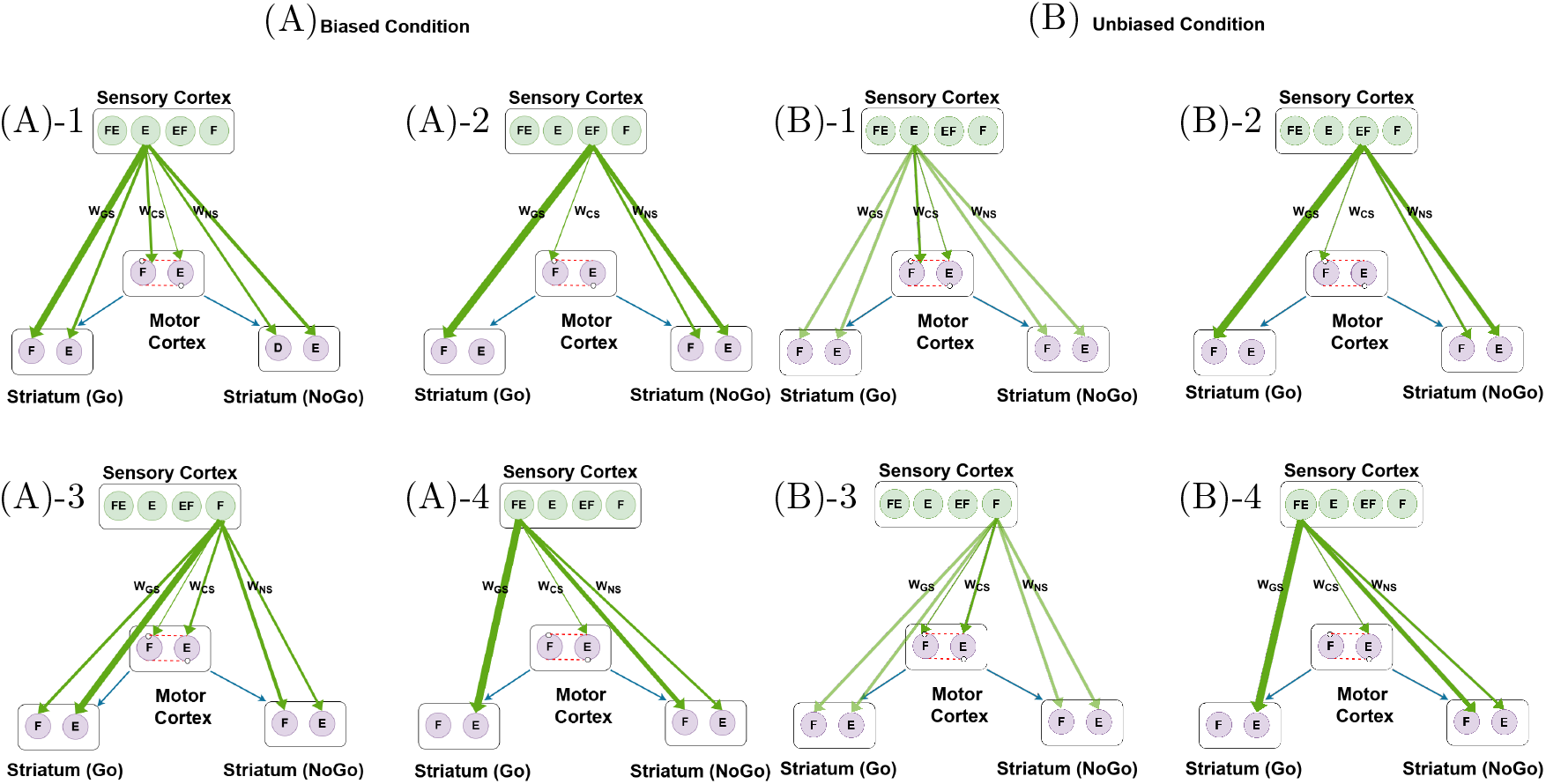
Representation of the synaptic connections in the *biased* and *unbiased* conditions, shown respectively in panel (A) and panel (B). Note that the two conditions differ primarily in how the *E* and *F* sensory signals project to the striatum. In the *biased* condition, the *E* and *F* sensory neurons project with unbalanced weights to the striatum. In the *unbiased* condition, the *E* and *F* sensory neurons project with identical synaptic weights to both the Go and NoGo neurons of the striatum.

Thus, the direct and indirect pathways are differentially engaged in promoting and suppressing motor actions. Simultaneous activation of the Go pathways for both E and F actions would imply the co-contraction of antagonistic muscles and a resulting increase in joint stiffness; this outcome is typically averted by the reciprocal inhibition implemented at the MC level. However, when the subthreshold accumulation of evidence between the cortical E and F neurons is closely matched, the hyperdirect pathway involving the subthalamic nucleus (STN) is recruited [33, 34]. This global activation provides a widespread inhibitory influence that effectively resets the cortical competition, thereby prolonging the DT. When multiple motor programs compete, the hyperdirect pathway acts as a dynamic brake, ensuring that only the ultimately selected program is executed by globally suppressing competing representations [20, 27]. This transient suppression lengthens the DT and, consequently, increases the ITI.

In summary, convergent signals from the sensory, motor, and premotor cortices induce a dynamic imbalance between the direct and indirect pathways of the CBGT loop. Specifically, the signals originating from the premotor cortex represent the motor intention, parameterized by 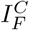 and 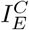 (see the fifth and seven-th rows of Eq. 7 in Method). The net imbalance between the Go and NoGo subnuclei is shaped by a combination of factors: the intensities of upstream inputs (including looped thalamocortical and sensori-cortical signals), synaptic weights modified via corticostriatal plasticity (*W*_*GS*_ and *W*_*NS*_ in Figure 5), and the level of dopaminergic neuromodulation, captured by the constant parameter *DA* as described by Eq. (11) in the Method section (see also the DA compartment in Figure 2). A well-calibrated imbalance between the Go and NoGo pathways ensures the smooth selection and execution of the desired movement, whereas an aberrant imbalance, including a loss of functionally relevant imbalance, manifests as motor deficits characteristic of Parkinson’s disease [20, 35].

### 2.4 Biased decision-making required for LRC

We first confirmed that a simple stochastic extension of the original Ursino model [20], incorporating additive motor noise into each neural mass population, failed to reproduce the characteristic LRC properties of human ITI time series. This limitation stems primarily from the absence of an explicit mechanical model of the effector. To address this, we integrated a mechanical index finger model into the computational CBGT framework to investigate how cycle-to-cycle action selection shapes the presence and magnitude of LRC in tapping variability. Guided by the finger’s equations of motion, the sensory cortex represents the time-delayed instantaneous mechanical state of the finger via the state vector *S* = [*S*_*FE*_, *S*_*E*_, *S*_*EF*_, *S*_*F*_ ].

A qualitative inspection of the simulated dynamics revealed that introducing a physical finger model alongside stochasticity is necessary but insufficient to generate the LRC properties of ITI time series. Based on a detailed inspection of the phase portrait of the finger dynamics (Figure 3, lower panels), we determined that state-dependent, biased competition for action selection between extension and flexion is the key mechanism generating tapping behavior with robust LRC properties under normal dopamine levels (*DA* = 0.5). Crucially, when this network-level action-selection competition was unbiased, the simulated ITIs failed to exhibit LRC, despite maintaining an oscillatory joint angle waveform consistent with standard finger tapping tasks.

This active competitive bias was introduced at the level of the sensoricorticostriatal projections—specifically, within the synaptic weights scaling the inputs from the sensory cortical E and F neurons to the Go/NoGo E and F populations. Figures 5(A) and (B) contrast the sensoricorticostriatal projection topologies that induce biased versus unbiased competitions. In Figure 5(A)-1, for instance, the projections from the sensory cortical E neuron to the Go and NoGo nodes are tuned to bias the system toward flexion. Consequently, when the delay-affected state of the finger is located in the region *S*_*E*_ (encoded as *S* = [0 1 0 0]), the flexion action is selected with a higher probability. Mechanistically, a large synaptic weight (*W*_*GS*_) from the sensory E neuron to the Go-F neuron facilitates the transition to flexion when the finger is extended, while a relatively large synaptic weight (*W*_*NS*_) from the sensory E neuron to the NoGo-E neuron concurrently suppresses the antagonistic extension action. Symmetrically, in Figure 5(A)-3, the imbalanced projections from the sensory F neuron—characterized by stronger connections to the Go-E and NoGo-F neurons alongside weaker connections to the Go-F and NoGo-E neurons—facilitate the selection of extension when the finger is flexed. Subtle, yet systematic, biased projections were likewise introduced for the transition phases between extension and flexion (state *S*_*EF*_, Figure 5(A)-2) and between flexion and extension (state *S*_*FE*_, Figure 5(A)-4). This active neural organization provides the necessary “synaptic bias” that drives fractal behavioral dynamics, distinct from passive mechanical dependencies.

Figure 6 shows typical examples of ITI time series with their corresponding thalamo-cortical signals and STN output obtained by simulating the model under three different conditions on sensoricortico-striatal projections as described in Table 1: namely, *biased* (panel A), *unbiased* (panel B), and *full competition* (panel C), under a standard dopamine level of *DA* = 0.5. In the biased condition (panel A), the ITI time series exhibits the structured low-frequency trends typically observed in well-organized, healthy motor behaviors. In contrast, the *unbiased* condition (panel B) displays a highly irregular pattern with reduced persistence of low-frequency trends. The persistence of such trends is completely lost in the *full competition* condition (panel C).

**Table 1.** Synaptic weight matrices used under three in silico conditions: *biased, unbiased*, and *full competition*.

| Condition |  | Synaptic Weights |  |  |  |  |  |  |  |
| --- | --- | --- | --- | --- | --- | --- | --- | --- | --- |
| *biased* | $W_{CS} = \begin{bmatrix} 0.0100 & 0.0018 & 0 & 0.0080 \\ 0 & 0.0080 & 0.0100 & 0.0018 \end{bmatrix}$ | $W_{GS} = \begin{bmatrix} 0.5000 & 0.1061 & 0 & 0.4243 \\ 0 & 0.4243 & 0.5000 & 0.1061 \end{bmatrix}$ | | | | $W_{NS} = \begin{bmatrix} 0.1000 & 0.2000 & 0.3000 & 0.1000 \\ 0.3000 & 0.1000 & 0.1000 & 0.2000 \end{bmatrix}$ | | | |
| *unbiased* | $W_{CS} = \begin{bmatrix} 0.0100 & 0.0018 & 0 & 0.0080 \\ 0 & 0.0080 & 0.0100 & 0.0018 \end{bmatrix}$ | $W_{GS} = \begin{bmatrix} 0.5000 & 0.1200 & 0 & 0.1200 \\ 0 & 0.1200 & 0.5000 & 0.1200 \end{bmatrix}$ | | | | $W_{NS} = \begin{bmatrix} 0.1000 & 0.1000 & 0.3000 & 0.1000 \\ 0.3000 & 0.1000 & 0.1000 & 0.1000 \end{bmatrix}$ | | | |
| *full competition* | $W_{CS} = \begin{bmatrix} 0.0100 & 0.0100 & 0 & 0.0100 \\ 0 & 0.0100 & 0.0100 & 0.0100 \end{bmatrix}$ | $W_{GS} = \begin{bmatrix} 0.5000 & 0.1200 & 0 & 0.1200 \\ 0 & 0.1200 & 0.5000 & 0.1200 \end{bmatrix}$ | | | | $W_{NS} = \begin{bmatrix} 0.1000 & 0.1000 & 0.3000 & 0.1000 \\ 0.3000 & 0.1000 & 0.1000 & 0.1000 \end{bmatrix}$ | | | |

**Fig 6.**
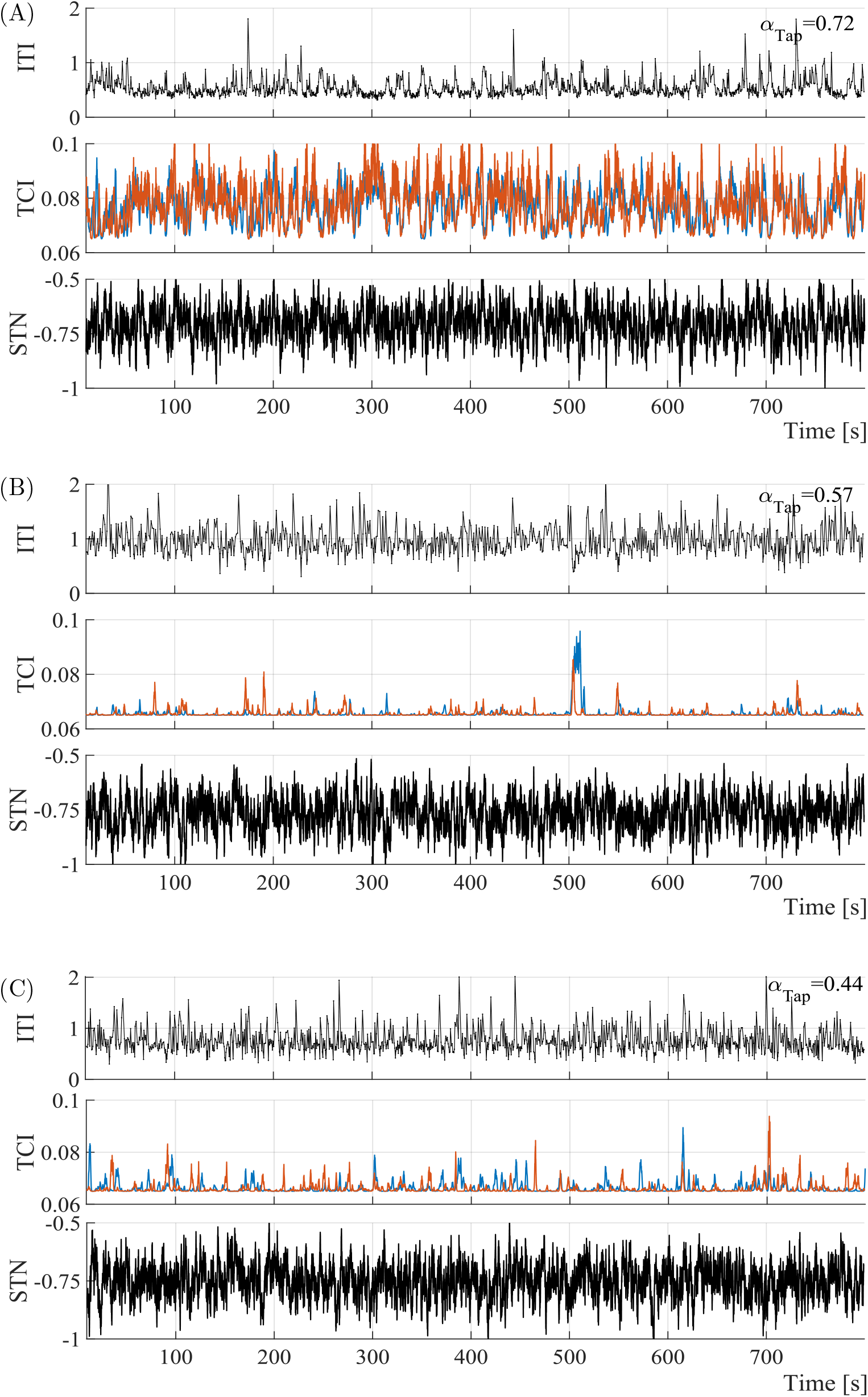
Long-range correlation analysis with standard DA=0.5. Panels (A), (B), and (C) show the time series for the ITI (intertap interval), TCI (thalamocortical input), and STN across the three different conditions. Blue and orange lines in TCI plots respectively indicate the activity of the F and E neural masses.

These time series, together with additional sample paths consisting of ten realizations of 800 seconds each, were analyzed using DFA. Figure 7 presents the DFA of the ITI and thalamocortical input (TCI) signals under the three conditions. As shown in Figure 7(A) for ITI, the mean scaling exponent in the *biased* condition was *α*_Tap_ = 0.70 *±* 0.04, which was significantly higher compared to both the *unbiased* condition (*α*_Tap_ = 0.50 *±* 0.06, *p* = 0.0003) and the *full competition* condition (*α*_Tap_ = 0.51 *±* 0.05, *p* = 0.0004). No significant differences were found between the *unbiased* and *full competition* conditions. Across all three conditions, the standard deviation of the fluctuations remained low up to a scale of 100 taps and increased slightly beyond that point, indicating limited sensitivity of the macroscopic ITI series to individual noisy realizations.

**Fig 7.**
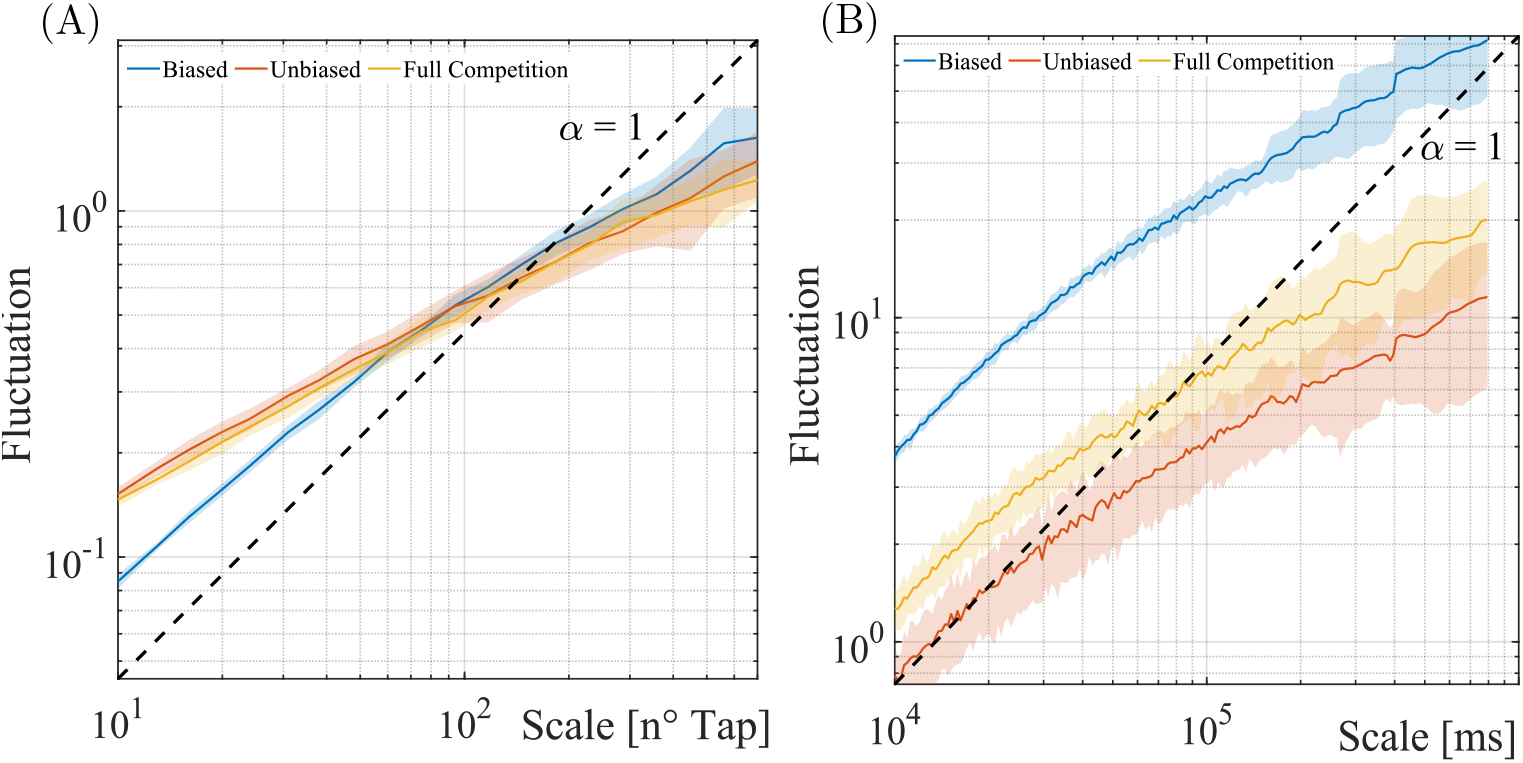
Fluctuation analysis of the ITI and TCI, shown in panels (A) and (B) respectively, for DA = 0.5, averaged across all simulations. The shaded areas represent the standard deviation of the fluctuations.

Figure 7(B) shows the average DFA curves for the thalamocortical signals. This analysis was performed on the thalamocortical signals for the cortical E and F neurons, excluding other driving inputs operating on the cortex. For consistency with the ITI analysis, the DFA for the thalamocortical signals covered the range from 10 to 500 seconds. The *biased* condition displayed stable fluctuations with minimal propagation in the standard deviation, indicating low sensitivity to noise. In contrast, the *unbiased* condition showed markedly higher variability, while the *full competition* condition presented an intermediate level of variability. The mean scaling exponent for the *biased* condition was *α*_TC_ = 0.67 *±* 0.05, compared to *α*_TC_ = 0.56 *±* 0.08 in the *unbiased* condition, and *α*_TC_ = 0.62 *±* 0.05 in the *full competition* condition. A statistically significant difference was confirmed between the *biased* and *unbiased* conditions (*p* = 0.0031), whereas no significant differences were observed between the *biased* and *full competition*, or between the *unbiased* and *full competition* conditions.

### 2.5 On the loss of complexity through dopamine depletion

A qualitative example of the effects of DA level reduction on the LRC of the ITI time series is shown in Figure 8. Comparisons among different DA levels are made with respect to the normal baseline (*DA* = 0.5) shown previously in Figure 6(A), for which an independent simulated sample path is reported in Figure 8(B), exhibiting a DFA *α*-value of 0.75.

**Fig 8.**
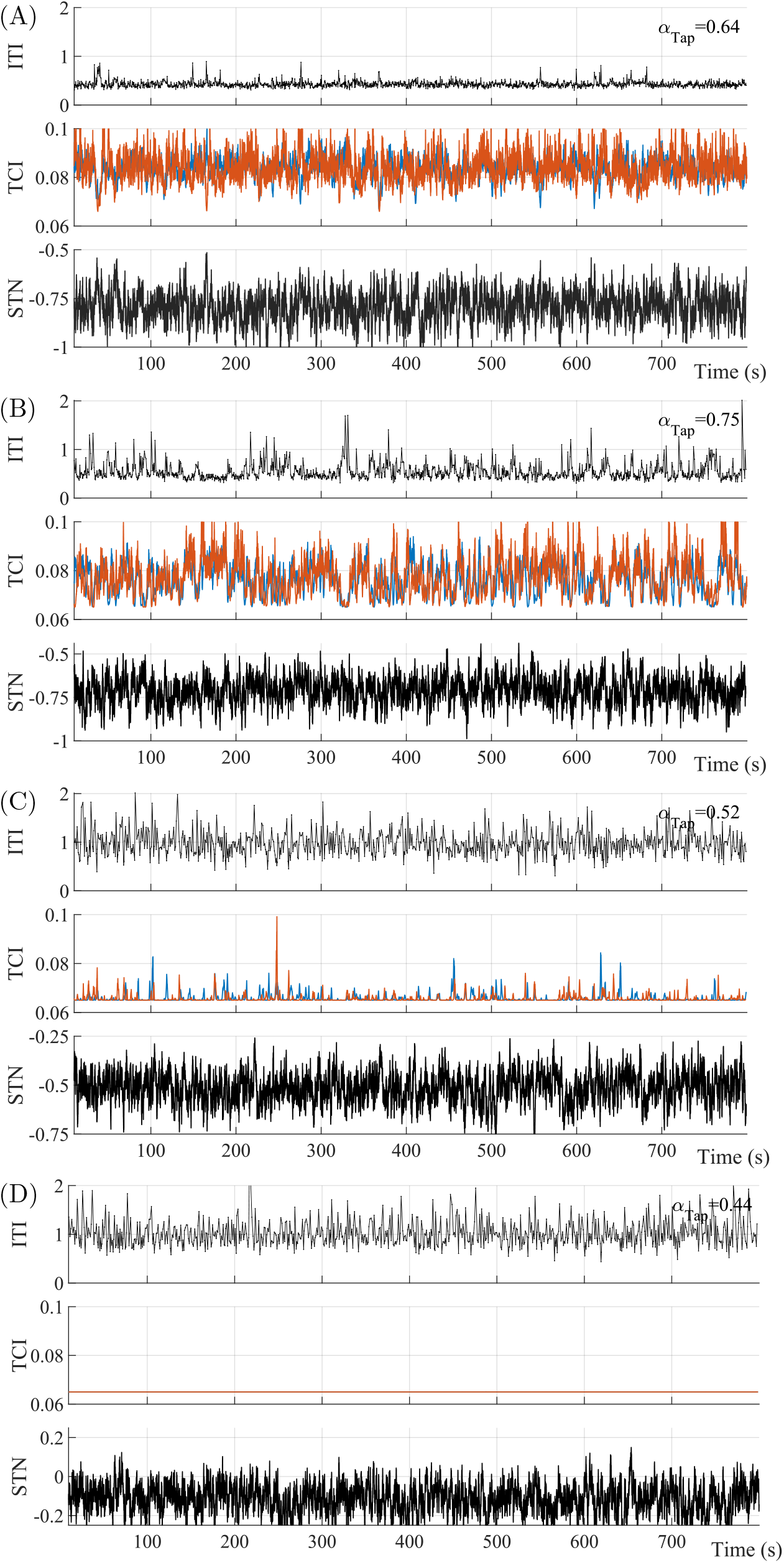
Examples of ITI, TCI, and STN activity simulated under the biased condition with decreasing dopamine levels, i.e., *DA* = 0.7, *DA* = 0.5, *DA* = 0.3, and *DA* = 0.1, are shown respectively in panels (A), (B), (C), and (D). Blue and orange lines in TCI plots respectively indicate the activity of the F and E neural masses.

As the DA level decreases (*DA* = 0.3 and *DA* = 0.1, Figure 8, panels C and D, respectively), both the ITI and STN outputs exhibit more random, white-noise-like behaviors, entirely stripping the time series of recognizable persistent trends. Conversely, at an elevated DA level (*DA* = 0.7, Figure 8, panel A), the ITI time series becomes less variable and highly repeatable, in which low-frequency structures are noticeably lost.

These qualitative observations were robustly confirmed by statistical analyses performed over the full set of simulations (ten realizations of 800 seconds for each condition). The bar plots of the scaling exponents as a function of the DA level for both ITIs and thalamocortical signals are presented in panels A and B of Figure 9. The results indicate that the *α*_Tap_ values obtained from our *in silico* experiments exhibit a progressive, monotonic decrease as DA levels reduce to 0.1. Specifically, the Kruskal-Wallis multiple comparison test revealed statistically significant differences between *DA* = 0.7 and *DA* = 0.1 (*p* = 0.0087), between *DA* = 0.5 and *DA* = 0.3 (*p* = 0.0004), and between *DA* = 0.5 and *DA* = 0.1 (*p <* 10^−4^). The highest mean value of *α*_Tap_ was 0.70 at *DA* = 0.5, while the lowest was 0.49 at *DA* = 0.1, mathematically demonstrating a transition from pink-noise-like persistent correlations to white-noise-like random behavior. This breakdown recapitulates a key qualitative feature of the “loss of complexity” documented in PD patients [36, 37].

**Fig 9.**
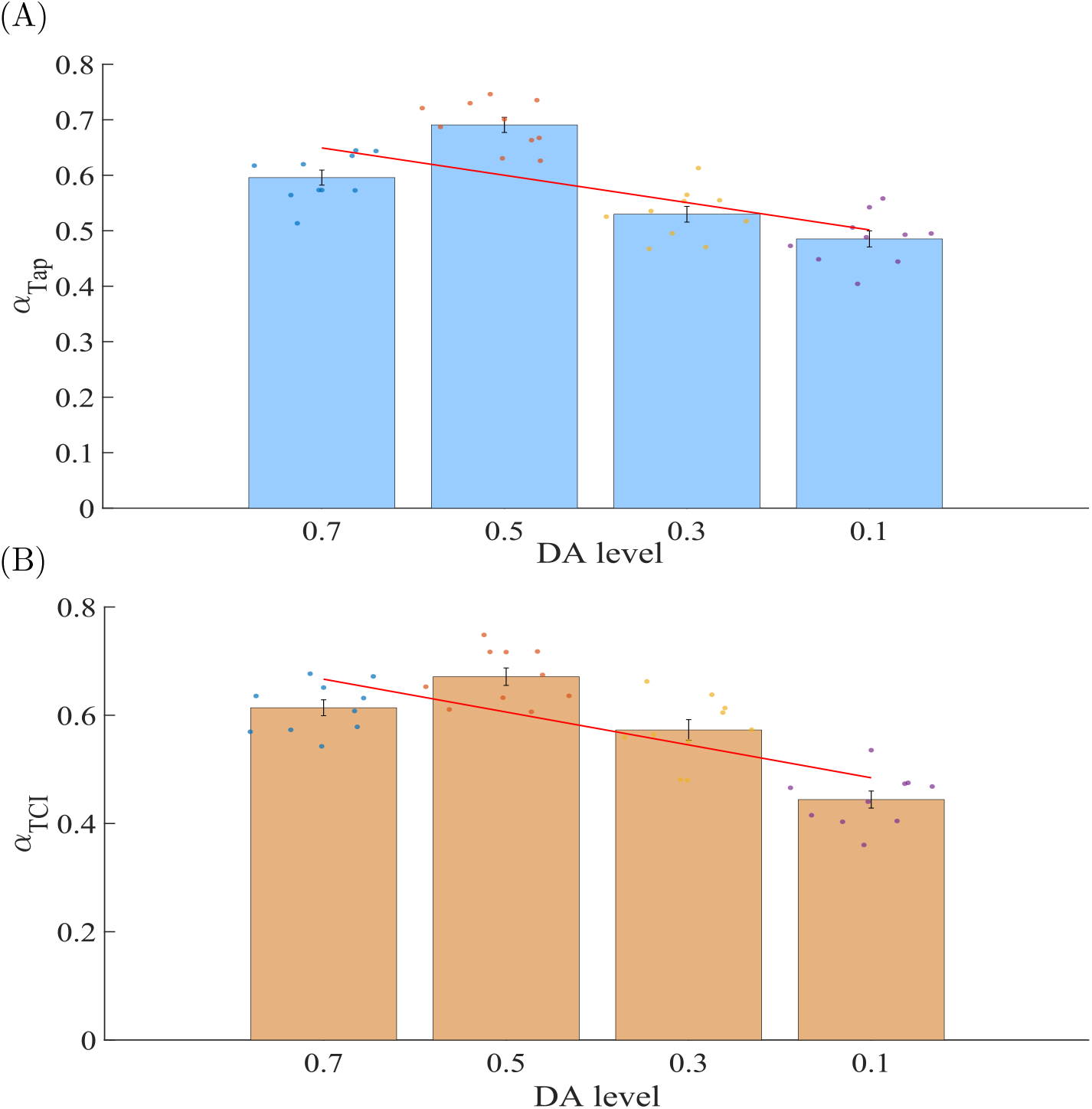
Bar plots of the average scaling exponents *α*_Tap_ and *α*_TIC_, corresponding to the inter-tap time series (panel A) and cortical input (panel B). Red lines highlight the linear trends computed across the averages of 10 in silico experiments for each condition.

The *α*_*TCI*_ values for the thalamocortical signals reported in panel B confirm a concurrent, significant decrease in long-range correlation properties when comparing *DA* = 0.7 to *DA* = 0.1 (*p* = 0.0017), *DA* = 0.5 to *DA* = 0.1 (*p <* 10^−4^), and *DA* = 0.3 to *DA* = 0.1 (*p* = 0.043). No statistically significant differences were observed in the other pairwise comparisons for the thalamocortical signals, underscoring that dopaminergic depletion disrupts macroscopic temporal complexity primarily by degrading the functional striatal selection gating.

## 3 Discussion

Neural mechanisms of decision-making have long been a central focus in the study of network functions within the CBGT loops [38, 39]. However, relatively few studies have addressed the embodied neural information processing associated with motor decision-making, where the tight integration of central neuromuscular control and peripheral musculoskeletal dynamics becomes indispensable [40]. Furthermore, automatic motor execution—such as finger tapping, quiet standing, and gait—has historically received less attention as behavioral paradigms that rely on continuous, subconscious decision-making. Recent neurophysiological evidence has begun to overturn this view, demonstrating that akin to volitional action selection, automatic motor control is strictly regulated by the CBGT circuitry, which dynamically determines when and which muscles the cortical motoneurons must activate or suppress [17–20].

The model proposed in this study bridges this gap by integrating the mechanical dynamics of a physical effector with a CBGT network responsible for the continuous decision-making that drives repetitive, rhythmic motor execution. This system operates under an intermittent control paradigm, driven by two distinct cortical neural masses: the cortical E neuron and the cortical F neuron. Within this framework, a WTA mechanism is implemented, grounded in drift-diffusion-like evidence accumulation and reciprocal cortical inhibition. Crucially, to generate rhythmic tapping patterns that match physiological time courses (see Figure 2), a robust WTA selection between flexion and extension must be resolved precisely when the finger state approaches the target extension angle *θ*_*E*_ (state *S*_*E*_) or flexion angle *θ*_*F*_ (state *S*_*F*_ ) in the phase plane. In our architecture, the degree of competition between these antagonistic cortical pathways is modulated by the synaptic strength of the sensoricorticostriatal projections, which defines the precise level of bias in decision-making (see Table 1).

We demonstrated that this biased decision-making—which dynamically facilitates the selection of flexor torque during the DT when the finger is extended, and vice versa—is a necessary and fundamental condition for generating long-range correlated ITI time series under healthy dopamine (*DA* = 0.5) levels. This persistent temporal structure was completely absent in both the unbiased and full competition conditions, which instead exhibited memoryless, white-noise-like ITI dynamics. These findings suggest that the presence of an asymmetrical synaptic bias within the striatal projection matrices (*W*_*GS*_ and *W*_*NS*_) actively steers the sequential execution of motor commands toward expressing fractal complexity. In other words, a functional imbalance in sensorimotor decision-making is vital to achieve smooth, robust transitions between physical states [41], directly giving rise to macroscopic LRC properties.

Furthermore, from a functional perspective, this active synaptic bias can be naturally interpreted as a structural representation of long-term motor learning and habituation. Throughout extensive trial-and-error motor practice, the neural circuit—particularly within the dorsolateral striatum (DLS) [42], which governs automaticity and habitual control—shapes its synaptic efficacy through activity-dependent plasticity. This learning process selectively strengthens the corticostriatal projection weights corresponding to the task-relevant [43], energetic-efficient state trajectory (i.e., biasing the striatal decision thresholds). Consequently, in healthy individuals, the neural controller does not solve a complex optimization problem in real time [44]; rather, it utilizes this pre-configured, experience-dependent “synaptic bias” to efficiently drive the phase point toward the deterministic limit cycle [45]. This mechanism allows the intermittent control framework to achieve rapid, robust motor execution with minimal real-time computational load [46, 47].

To clarify the mechanistic role of this synaptic bias in generating LRC, it is highly insightful to contrast our findings with recent computational frameworks on intermittent postural control during quiet standing [48]. Human upright postural sway exhibits fractal LRCs remarkably similar to those observed in healthy finger tapping intervals. In the postural control framework, the intermittent control paradigm is similarly implemented via a CBGT loop [29], where overall system stability and LRC emerge from a continuous switching between active feedback control (control-ON) and the passive mechanical dynamics of an inverted pendulum (control-OFF) [48]. Crucially, however, in the postural CBGT model, the active control is turned off during every DT while the network engages in an unbiased competition between antagonistic motor commands (dorsiflexion vs. plantarflexion) [29]. During these intermittent control-OFF periods, the postural system exploits an intrinsic “mechanical bias” provided entirely by the body’s passive physics: because the passive state of the pendulum, under the gravitational toppling torque, lies near the stable manifold of a saddle-type unstable equilibrium [15, 46], the passive mechanics in the vicinity of the stable manifold naturally guide the body back toward the upright vertical position. This combination of a neurally unbiased DT and a physical, passive mechanical bias creates long-period orbits near a homoclinic trajectory, which, when perturbed by stochastic noise, naturally emerges as behavioral LRC.

In sharp contrast, the volitional finger tapping task investigated in the present study entirely lacks such an intrinsic, stabilizing mechanical bias. Although the finger’s physical parameters (such as inertial and passive viscoelastic forces) are explicitly incorporated into our equations of motion, there is no corresponding stable manifold to naturally or automatically guide the finger between extension and flexion while the CBGT network is actively resolving its choice during the intermittent control-OFF period [49]. Consequently, if the competition within the current finger-tapping CBGT network were completely unbiased, the system would select extension and flexion with almost equal probability at the kinematic reversal points (noting a small bias due to inertia of the finger movement), immediately disrupting rhythmic execution and collapsing into random variability. Therefore, introducing an explicit “synaptic bias” into the sensoricorticostriatal projections [50], which prioritizes the selection of flexion when the finger is extended, and vice versa—becomes a necessary, mandatory condition unique to the finger-tapping model to achieve smooth state transitions and sustain temporal complexity.

This discrepancy in the neural competition dynamics between finger tapping and postural control does not diminish the generality of our model; rather, it highlights a fundamental, unifying principle of embodied neural control. The contrast between these two CBGT paradigms demonstrates that a specific neural bias is not a universal, hardcoded prerequisite for LRCs in motor behavior. Instead, the emergence of temporal complexity is determined by a complementary interplay between the central nervous system and peripheral musculoskeletal dynamics [51]: where nature provides an intrinsic mechanical bias (as in quiet standing), the neural controller can remain unbiased; where such mechanical assistance is absent (as in volitional finger tapping), the neural controller must actively introduce a structured synaptic bias to smoothly steer behavioral transitions. Ultimately, our results suggest that temporal complexity in motor output is a strictly emergent property arising not from the brain or biomechanics in isolation, but from the tightly coupled, closed-loop interaction between both.

A related principle has also emerged from studies of human walking, where interlimb coordination exhibits a dead zone around the antiphase state, such that corrective control is effectively withheld until phase deviations exceed a threshold [52]. In addition, computational studies of bipedal gait have shown that intermittent control, together with event-driven phase resetting, can generate 1*/f* -like gait-cycle variability near the edge of stability [53]. Together, these observations point to a broader principle whereby temporal complexity emerges when motor control deliberately leaves periods of unconstrained dynamics, while sparse state- or event-dependent interventions bias the system toward appropriate transitions; the present CBGT architecture may implicitly embody both features through its control-free decision intervals and sensoricorticostriatal synaptic bias.

Although the present model does not implement an explicit online learning rule [54], the biased condition validated here may reflect a steady-state synaptic organization acquired through motor learning [55], and the long-term consolidation of automated movements [56]. This interpretation aligns with neurophysiological evidence identifying the dorsolateral striatum (DLS) as a primary locus for habit formation [57], automaticity, and motor skill consolidation, while the tail of the striatum acts as a functional filter for sensorimotor integration [58]. Within this framework, the sensory-driven modulation of the Go/NoGo populations in our proposed model [59], implemented through *W*_*GS*_ and *W*_*NS*_, represents a simplified computational analogue of DLS gating. Consequently, the emergence of LRCs in the ITI time series under the biased condition can be interpreted as the direct outcome of a stabilized, learned synaptic topology that supports optimized motor sequencing and robust sensorimotor feedback, consistent with the principles of motor automaticity.

Conversely, the loss of motor complexity in Parkinsonism may reflect pathological remodeling of corticos triatal transmission, which disrupts the functional balance between direct and indirect pathways and degrades sensorimotor integration [60, 61, 61–64]. In our model, the unbiased and full-competition conditions provide simplified computational analogues of such disrupted synaptic organization, in which the loss of appropriate sensoricorticostriatal bias impairs state-dependent action selection and yields disorganized, memoryless motor output [65].

By analyzing the characteristics of the internal thalamocortical signals, namely TCI, we confirmed that across the longer time scales analyzed, the internal network fluctuations exhibit markedly different behaviors among the three conditions (Figure 6). However, the scaling exponent showed statistically significant differences only between the biased and unbiased conditions. To unravel this nuance, we examined the specific role of the STN in the hyperdirect pathway [66]. Circuit-wise, the STN monitors the overall energy of MC activity and acts as a global brake to mitigate high levels of conflict during intense competition between opposing motor programs [67].

The mean power of simulated STN activity (see Table 2) demonstrates that the hyperdirect pathway effectively reflects the level of underlying network conflict. The unbiased condition, which exhibited the highest sensitivity to noise realizations, was characterized by a scaling exponent similar to that of the full competition condition (*α* ≈ 0.5). However, its STN activity was the most prominent across all conditions. Because the unbiased weights for *W*_*GS*_ and *W*_*NS*_ maximize the competition within the basal ganglia subnuclei, they trigger an immense level of conflict and network instability. The STN attempts to mitigate this conflict by increasing global inhibition via the indirect pathway [67], as evidenced by the significantly greater mean STN power in the unbiased condition compared to both the biased and full competition baselines (see Table 2). This intense hyperdirect engagement ultimately drives the cortical WTA mechanism into highly erratic, random behavior. In contrast, in the full competition condition, the absence of synaptic biases in both the sensoricortex-to-cortex (*W*_*CS*_) and sensoricortex-to-striatum pathways statistically flattens the conflict between the MC and BG circuitry. This yields an intermediate mean STN power. Thus, while the scaling exponents of the TC signals in the unbiased and full competition conditions are similarly non-persistent at long time scales, the underlying hyperdirect conflict-handling mechanism differs fundamentally, with the unbiased condition exhibiting the highest level of network friction and hyperdirect pathway recruitment.

**Table 2.**
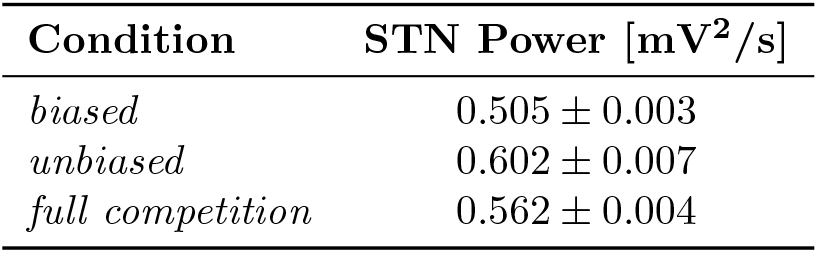
Mean and standard deviation of STN signal power under the three different conditions. Statistical analysis reveals significant differences in all the comparisons with *p <* 0.05.

A crucial contribution of this study concerns the progressive deterioration of motor control and the associated loss of LRC properties under conditions of altered dopamine levels. While classic basal ganglia models have demonstrated that dopamine depletion leads to bradykinesia and amplitude reduction, our *in silico* framework proves that by combining neural masses with explicit biomechanical execution and sensory coding, we can capture the precise degradation of temporal complexity as a continuous function of DA levels (see Figure 9). Our statistical analyses revealed a monotonic decrease in *α*_Tap_ as DA levels fell from a healthy baseline (*DA* = 0.5) to severe depletion (*DA* = 0.1), driving a clear transition from adaptable, pink-noise-like persistent correlations to rigid, white-noise-like random behaviors. This breakdown faithfully recapitulates a key qualitative feature of the clinical “loss of complexity” observed in PD.

Consequently, the neuromechanical model developed here provides a powerful computational engine for investigating how graded neuromodulation shapes macroscopic behavioral dynamics. For clinical translation, statistical inference techniques such as Approximate Bayesian Computation can be deployed to map patient-specific finger-tapping data to individual model parameters [15]. This allows for the creation of high-fidelity “digital twins” of individual patients.

We acknowledge that the present study is not without limitations. Although parameter tuning was strictly guided by established literature values, incorporating online synaptic learning rules (such as dopamine-dependent) could provide deeper insights into how the CBGT circuitry actively adapts to progressive dopaminergic cell loss over time [60]. However, this lies beyond the current scope of this study, which aims to establish the baseline neuromechanical framework and demonstrate how sensory-driven synaptic biasing acts as the primary driver of LRC properties.

Advanced computational models are increasingly valuable complementary tools for understanding motor disorders and developing patient-specific approaches to in silico neurology [68, 69]. In this context, the ability of a neuromechanical model to reproduce physiologically plausible LRC—and their breakdown through explicit parametric alterations—may provide an important mechanistic foundation for future digital-twin applications. By integrating finger biomechanics with CBGT decision-making, the present framework provides such a foundation while linking neural circuit alterations to macroscopic temporal complexity. Future work will incorporate reinforcement learning-based synaptic adaptation under varying DA conditions [54], allowing *W*_*CS*_, *W*_*GS*_, and *W*_*NS*_ to evolve dynamically and enabling investigation of dopamine-dependent compensatory mechanisms.

## 4 Conclusion

In conclusion, we developed an embodied computational model integrating CBGT neural dynamics with finger biomechanics under intermittent control. The model demonstrates that LRC in finger tapping can emerge from state-dependent sensoricorticostriatal bias interacting with physical execution, contrasting with the mechanical bias previously identified in intermittent postural control. Dopamine depletion progressively disrupts this temporal organization, recapitulating a key qualitative feature of the loss of motor complexity observed in PD. Together, these findings support a broader view in which physiological temporal complexity emerges from closed-loop interactions between neural decision-making and body dynamics rather than from either component alone. The framework may therefore provide a mechanistic foundation for future patient-specific modeling of motor dysfunction.

## 5 Methods

### 5.1 Human Finger Tapping Experiment

This study was conducted in accordance with the Declaration of Helsinki and approved by the Ethics Committee of the Graduate School of Informatics, Kyoto University. Written informed consent was obtained from the participant prior to the experiment.

The participant, a right-handed healthy young adult, performed a rhythmic finger tapping task using the right index finger. The task was initiated with the right hand placed in a comfortable resting position in front of the abdomen. The experiment consisted of two consecutive phases: a synchronization phase and a continuation phase. During the initial synchronization phase, auditory metronome cues were presented at a frequency of 1.8 Hz for approximately the first 10 taps to establish a stable time baseline. Following the cessation of the metronome cues, the participant continued self-paced finger tapping based on their internal rhythm for 250 seconds (continuation phase).

Kinematic data of the repetitive finger flexion and extension were recorded using a compact tri-axial accelerometer (HapLog, Tech Gihan Co., Ltd., Kyoto, Japan) attached to the index fingertip. Inter-tap intervals (ITIs) were defined as the time duration between consecutive extension acceleration peaks.

### 5.2 Detrended Fluctuation Analysis (DFA)

To evaluate the long-range correlation (LRC) properties of the ITI time series obtained from both the human experiment and the computational model simulations, we performed DFA, which is a modified random-walk analysis designed to quantify self-similarity and scale-invariant long-range correlations while mitigating non-stationarities and local trends.

Given an ITI time series *x*_*i*_ (*i* = 1, 2, …, *N*, where *N* is the total number of taps), the series is first integrated:

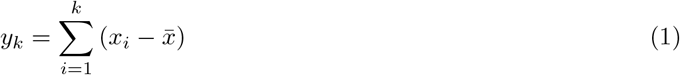

where 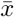 denotes the mean value of the time series. The integrated series *y*_*k*_ is then divided into non-overlapping windows of length *n*. Within each window, a local trend *y*_*n,k*_ is estimated using a second-order polynomial fit (least-squares method). The root-mean-square fluctuation *F* (*n*) of the detrended series is calculated as:

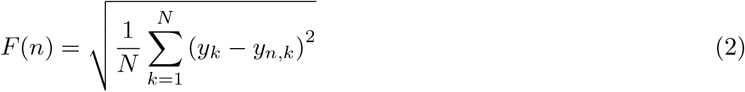

The fluctuation function *F* (*n*) typically exhibits a power-law relationship with the window size *n*, represented as *F* (*n*) ∝ *n*^*α*^. The scaling exponent *α* is determined by calculating the slope of the linear regression fit between log_10_ *F* (*n*) and log_10_ *n*.

An exponent of *α* = 0.5 indicates uncorrelated white noise, whereas *α >* 0.5 indicates positive persistence (long-range correlations), where larger intervals tend to be followed by larger intervals. In this study, local trends were evaluated using second-order polynomial fitting (DFA2) for window sizes spanning 6–500 cycles, and the scaling exponent *α* was obtained via linear regression over the scaling range of *n* ∈ [30, 200] cycles.

### 5.3 Overview of the CBGT Neural Network

The CBGT neural network used to simulate the motor decision-making was based on the models proposed in [20, 27]. However, major modifications were made to incorporate a sensory representation of finger mechanics and to realize control of finger mechanical dynamics based on the output of cortical motor neurons. This section provides a mathematical description of the model, highlighting its structure and the novel aspects introduced. After brief overview of the model, detailed equations and a set of the tuned parameters are presented to ensure completeness and reproducibility of the study.

Figure 2 shows the overall structure of the CBGT model. MC is composed of two Morris–Lecar neurons acting as a stochastically coupled oscillator [32, 70], where one neuron is responsible for selecting the E action and the other for the F action. Instead of employing a if-then-rule-based WTA mechanism with waiting times [20], WTA behavior here emerges dynamically through a competition between two cortical neurons based on a drift-diffusion-like model. That is, each neuron receives signals through the thalamocortical projections and those from the sensory cortex. The two MC neurons are reciprocally inhibited via voltage-dependent synaptic conductance, which promote competition and enable strong coupling of the oscillator [32]. A first novel aspect regards the use of cortical neural activity to implement intermittent control law for generating finger movement (see Figure 2 ) Indeed, assuming that 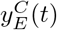 and 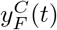 are respectively the output signal of the cortical E and F neurons, an active torque *τ*_*a*_(*t*) selection rule was framed as follows:

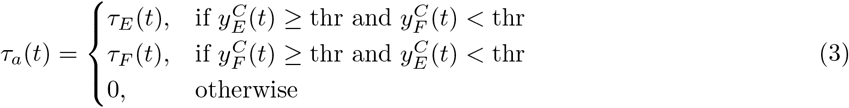

where *τ*_*E*_(*t*) and *τ*_*F*_ (*t*) implement mechanical feedback to steer the finger toward the desired extended or flexed position, respectively (see Method). The variable `thr` represents a decision threshold on the cortical signal (see Table 6), used to reflect the outcome of the WTA mechanism occurring at the MC level onto the mechanical response of the finger.

Although torque selection is performed at the cortical level, each selection is made through the downstream decision-making network. Following the approach in [20, 27], the model incorporates the direct, indirect, and hyperdirect pathways of BG, whose components are listed in Table 3 and depicted in Figure 2. These pathways are reconnected with the cortex through a positive feedback loop involving the thalamus, which plays a critical role in generating the instabilities characteristic of WTA circuits [20]. As a result, the selected torque corresponds to one of three functional states of the BG: competition between E and F actions, yielding zero torque; selection of E; or selection of F, each determined by the dominance of the Go pathway over the NoGo pathway for the respective action. When strong competition occurs in the cortex, the hyperdirect pathway inhibits thalamic activity, helping to preserve orderly alternation between E and F neurons in the cortical neuron. However, the model allows for consecutive cortical activations of E and F states. This variability, filtered by the mechanics of the finger, can produce physiologically plausible complexity in the resulting sway behavior, offering a potential source of long-range correlations in motor output.

**Table 3.** Functional components of BG pathways employed in the model. The table reports the nomenclature and recalls the structure of each pathway [3, 20, 27].

| Pathway | Components | Functional Role |
| --- | --- | --- |
| Direct | Motor Cortex (MC), Go part of the striatum (Go), Globus Pallidus pars interna (GPi), Thalamus (T) | Facilitates action selection by disinhibiting the thalamus, allowing the cortex to execute the chosen motor command. |
| Indirect | Motor Cortex (MC), NoGo part of the striatum (NoGo), Globus Pallidus pars externa (GPe), Subthalamic Nucleus (STN), GPi, Thalamus (T) | Suppresses unwanted or competing motor programs by increasing inhibition on the thalamus through additional relays. |
| Hyperdirect | Motor Cortex (MC), Subthalamic Nucleus (STN), GPi, Thalamus (T) | Provides fast, global inhibition of all motor programs via the STN to resolve conflicts or prevent premature action selection. |

### 5.4 Finger Mechanics

The finger sway movement was modeled in the sagittal plane, using a pendulum mechanism with the base of the pendulum fixed at the metacarpophalangeal (MCP) joint. This joint is modeled as a revolute joint, while the proximal interphalangeal and distal interphalangeal joints are assumed to remain fixed during finger movement, as shown in Figure 2. Given the small inertia of the finger, the gravitational component can be neglected, and the motion of the pendulum can be well approximated by a second-order linear differential equation, as described in [71]:

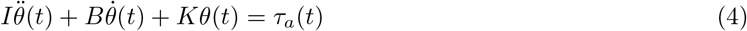

Where *I, B*, and *K* represent, respectively, the moment of inertia, the equivalent torsional damper, and the torsional spring at the MCP joint [71], while *θ*(*t*) denotes the finger sway angle with respect to the horizontal axis in the sagittal plane. As discussed in Section 5.3, the torque *τ*_*a*_(*t*) is intermittently selected among three possible values as defined in Equation (3). When the E or F neuron in the MC wins the competition, the active torque switches from 0 to *τ*_*E*_(*t*) or *τ*_*F*_ (*t*), respectively:

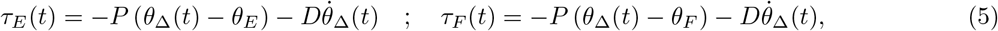

where, *θ*_Δ_(*t*) and 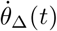 represent the angular position and angular velocity delayed by a time Δ, which accounts for afferent and efferent transmission delays in the peripheral nervous system. *P* and *D* are proportional and derivative control gains implementing the neuromuscular feedback. The variables *θ*_*E*_ and *θ*_*F*_ define the desired angular positions corresponding to the final extension and flexion finger configurations. These two values cause the mechanical system to oscillate around the two target angles. The parameters used in the finger dynamics simulation are summarized in Table 4.

**Table 4.** Model parameters used for simulating the finger sway dynamics.

| Parameter | Value |
| --- | --- |
| $K$ | 0.001 [Nm/rad] |
| $B$ | 0.0005 [Nms/rad] |
| $I$ | $2.5 \times 10^{-5}$ [kg·m<sup>2</sup>] |
| $P$ | 0.008 [Nm/rad] |
| $D$ | 0.0005 [Nms/rad] |
| $\theta_E$ | 0.5236 [rad] |
| $\theta_F$ | -0.5236 [rad] |
| $\Delta$ | 0.06 [s] |

### 5.5 Sensory information coding

The angular displacement and velocity were used as state variables for the sensory cortex, which is responsible for encoding this information and making it available to both the MC and the Go and NoGo parts of the striatum, in order to contribute to the decision-making process (see Figure 2). To achieve this, a coding rule based on the phase portrait of the delayed state of the pendulum was designed by partitioning the phase space into four regions, as illustrated in Figure 4. The zones *S*_*E*_ and *S*_*F*_ represent regions in which the sensory cortex interprets the finger as being close to the desired positions *θ*_*E*_ and *θ*_*F*_, respectively. In contrast, *S*_*FE*_ and *S*_*EF*_ are transitional zones indicating that the finger is moving from F to E or from E to F. Each zone corresponds to an input neuron in the sensory cortex that switches between binary states (0 or 1), as shown in Figure 2. The resulting coding rule allows the conversion of mechanical state information of the finger into a neural representation in the sensory cortex, modeled as follows:

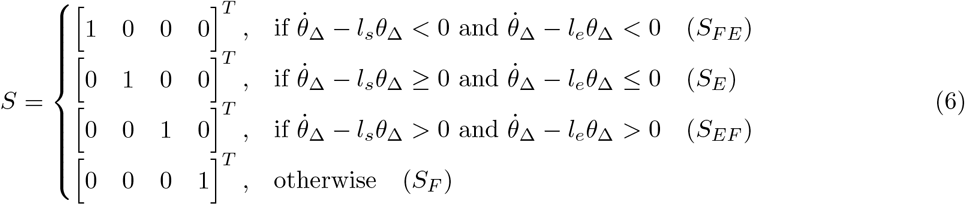

The slopes *l*_*s*_ and *l*_*e*_ were set to 50 s^−1^ and −10 s^−1^, respectively. Such input signal *S* was injected into the network using different weighting matrices, depending on the *in silico* experiments, as described in the following section.

### 5.6 *in silico* experiments

A first *in silico* experiment conducted with the model aimed to verify its capability to exhibit long-range correlation properties in the inter-tap time intervals of the finger. To do this, synaptic weight values consistent with non-pathological conditions, as described in [20], were considered for the BG circuitry and reported in the Method (Table 5). However, to accommodate the new formulation of the sensory input to the Morris–Lecar cortical oscillator and the Go and NoGo populations, three new synaptic matrices were tuned. Specifically, to reproduce long-range correlation properties in the inter-tap time series, it was hypothesized that such synapsis should be biased to excite the cortical F neuron when the finger is in the E state, and vice versa. The corresponding synaptic weights for sensory input to the Go population (*W*_*GS*_), to the NoGo population (*W*_*NS*_), and to the cortex (*W*_*CS*_) were reported in Table 1 under the *biased*, condition.

**Table 5.** Parameters of the basal BG and associated compartments used in the model. Abbreviations: C = Motor Cortex, G = Go, N = NoGo, E = GPe, I = GPi, STN = Subthalamic Nucleus, T = Thalamus, H = Cholinergic Interneurons.

| Parameter | Structure | Meaning | Value |
| --- | --- | --- | --- |
| $\tau$ | Scalar | Common neural time constant | 0.5 |
| $a$ | Scalar | Sigmoid slope for BG neurons | 4 |
| $u_0$ | Scalar | Sigmoid center for BG neurons | 1 |
| $\delta^G$ | Scalar | Go threshold | 0.35 |
| $I^E$ | Scalar | Bias input to GPe | 1 |
| $I^I$ | Scalar | Bias input to GPi | 3 |
| $I^H$ | Scalar | Bias input to cholinergic interneurons | 1 |
| $\gamma$ | Scalar | DA modulation for Go neurons | 0.75 |
| $\beta$ | Scalar | DA modulation for NoGo neurons | -1 |
| $\eta$ | Scalar | DA modulation for cholinergic interneurons | -0.5 |
| $k_E$ | Scalar | STN energy/conflict gain | 0.4 |
| $W_{GC}$ | Diagonal matrix | Synaptic weight from C to G | $w_{ii}^{GC} = 0.580$ |
| $W_{NC}$ | Diagonal matrix | Synaptic weight from C to N | $w_{ii}^{NC} = 0.399$ |
| $W_{EN}$ | Diagonal matrix | Synaptic weight from N to GPe | $w_{ii}^{EN} = -2.2$ |
| $W_{IE}$ | Diagonal matrix | Synaptic weight from GPe to GPi | $w_{ii}^{IE} = -3$ |
| $W_{IG}$ | Diagonal matrix | Synaptic weight from G to GPi | $w_{ii}^{IG} = -36$ |
| $W_{TC}$ | Diagonal matrix | Synaptic weight from C to Thalamus | $w_{ii}^{TC} = 0.7$ |
| $W_{TI}$ | Diagonal matrix | Synaptic weight from GPi to Thalamus | $w_{ii}^{TI} = -3$ |
| $w^{ESTN}$ | Scalar | Synaptic weight from STN to GPe | 1 |
| $w^{ISTN}$ | Scalar | Synaptic weight from STN to GPi | 30 |
| $W^{STNE}$ | Row vector | Synaptic weight from GPe to STN | $w_j^{STNE} = -1$ |
| $w^{GH}$ | Scalar | Synaptic weight from H to Go | -1 |
| $w^{NH}$ | Scalar | Synaptic weight from H to NoGo | 1 |

In the second experiment, referred to as the *unbiased* condition, the matrix *W*_*CS*_ was kept unchanged from the first experiment, while a higher level of competition was introduced in the direct and indirect pathways. This was done by assigning equal values in *W*_*GS*_ and *W*_*NS*_ when the finger is sensed in the E and F regions, respectively (see Table 1). It was hypothesized that this increased competition may lead to local disruptions in the temporal ordering of motor command selection, i.e., a more random behavior of the WTA mechanism in the MC, which could in turn, reduce the long-range correlation properties observed in the inter-tap interval time series. A third experiment was designed to evaluate whether long-range correlations in the inter-tap interval time series may persist when an unbiased synaptic configuration is also applied to the cortical weights *W*_*CS*_. This condition, labeled as the *full competition* condition (see Table 1), assumed that sensory information about the state of the finger has minimal influence on the decision-making process. In this case, the weights in *W*_*CS*_ for the E and F regions were set equal, while the same competition levels in *W*_*GS*_ and *W*_*NS*_ from the *unbiased* condition were maintained. As a result, the direct, indirect, and hyperdirect pathways may produce less structured responses, rendering the selection of E or F actions more stochastic.

An additional experiment was set to investigate the detrimental role of DA level reduction in long-range correlation properties. Indeed, assuming *biased* synaptic weights, DA was varied across four values, *DA* ∈ *{* 0.7, 0.5, 0.3, 0.1}, representing a decreasing level from normal to conditions typically associated with Parkinsonian states [36].

Under this experimental framework, ten simulations of 800 s each were run for every experimental condition and DA level, generating time series data for cortical, BG, and finger dynamics. It is worth noting that the noise realizations acting on the system (see Method) were characterized by identical statistical properties but employed different random seeds, thereby introducing variability in the temporal evolution of both neural and mechanical activity. A long-range correlation analysis was performed on two different types of time series. The first involved the inter-tap interval time series, computed as the sequence of time intervals between successive peak points in the finger sway corresponding to the U state. The second analysis focused on the input signal to the cortex. If the inter-tap interval time series exhibited long-range correlation properties, these would be expected to originate from the decision-making system itself and should be reflected in the closed-loop dynamics of the entire neural circuitry, particularly in the cortical input influencing WTA mechanism. For both analyses, DFA was applied, and the scaling exponent *α* was computed, i.e. *α*_Tap_ and *α*_TCI_, respectively for inter-tap time interval and input signal to the cortex. Non-parametric Kruskal-Wallis tests were conducted on the *α* values obtained under each DA level and significance threshold was set at 0.05.

### 5.7 Mathematical Model of the Neural Circuitry

In this subsection, a mathematical description of the neural circuitry as well as the parameters setting for implementing the neural model has been reported in subsections.

#### Motor Cortex

The MC was modeled using a Morris–Lecar coupled oscillator [32]. Specifically, the dynamics of this system, consisting of E and F neurons connected via event-driven double-exponential synapses, was defined as follows,

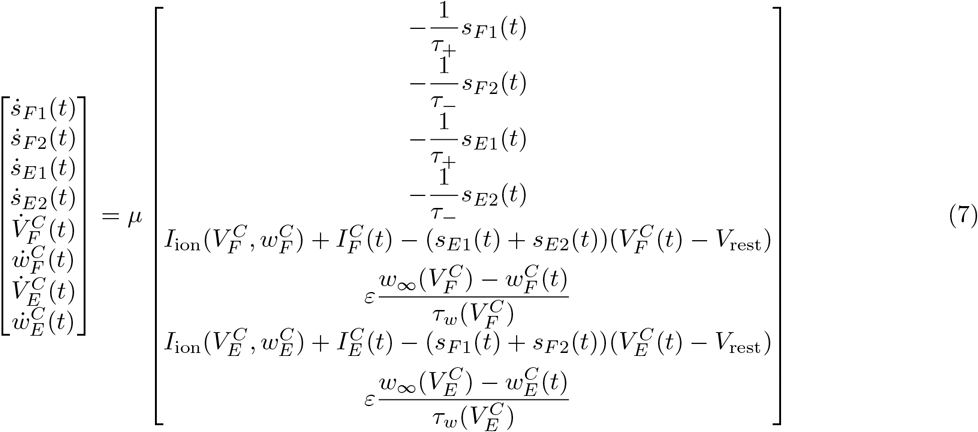

where *s*_*F* 1_, *s*_*F* 2_, *s*_*E*1_, *s*_*E*2_, are co-states used to implement the event-driven double-exponential synapses. These variables decay exponentially with time constants *τ*_+_ and *τ*_−_ and are instantaneously updated when the presynaptic neuron voltage crosses the threshold *V*_thr_ (see eq. 8). 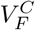 and 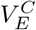 represent, respectively, the membrane potentials of the flexion (*F* ) and extension (*E*) cortical neurons, while 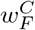 and 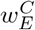 are the corresponding recovery variables controlling the activation of the potassium current in the Morris-Lecar model [32, 72]. As mentioned the decay exponential was implemented with an updating rule as reported in eq. (8), here *t*^−^ and *t*^+^ represent respectively the previous and the current step of integration.

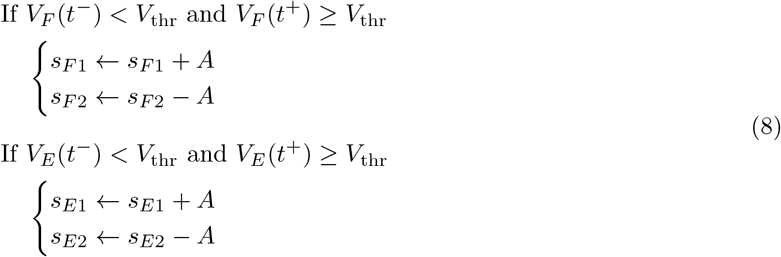

Further details regarding the functions and parameters employed are reported in Table 6. An exception is made for the two terms 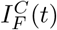 and 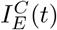, which represent the input currents to the cortical neurons. These currents depend on the interaction with the BG circuitry, as illustrated in Figure 2 and described in the next section. The membrane potentials *V*_*F*_ and *V*_*E*_ are mapped to the neuronal output using a sigmoid function of the form

**Table 6.** Functions, and parameters used in the Morris–Lecar coupled oscillator to simulate the motor cortex. Values are reported without units for simplicity.

| Function / Parameter | Expression | Value |
| --- | --- | --- |
| Ionic current | $I_{\text{ion}}(V, w) = -g_{\text{Ca}} m_{\infty}(V)(V - E_{\text{Ca}}) - g_K w(V - E_K) - g_L(V - E_L)$ | – |
| Steady-state activation | $m_{\infty}(V) = \frac{1}{2} \left[ 1 + \tanh \frac{V+0.01}{0.15} \right]$ | – |
| Steady-state recovery | $w_{\infty}(V) = \frac{1}{2} \left[ 1 + \tanh \frac{V-0.1}{0.145} \right]$ | – |
| Recovery time constant | $\tau_w(V) = \frac{1}{\cosh \frac{V-0.1}{0.290}}$ | – |
| $g_{\text{Ca}}$ | – | 1.33 |
| $g_K$ | – | 2.0 |
| $g_L$ | – | 0.5 |
| $E_{\text{Ca}}$ | – | 1.0 |
| $E_K$ | – | -0.7 |
| $E_L$ | – | -0.5 |
| $V_{\text{rest}}$ | – | -0.4 |
| $\varepsilon$ | – | 0.333 |
| $\tau_+$ | – | 0.08 |
| $\tau_-$ | – | 0.03 |
| $V_{\text{thr}}$ | – | 0.2 |
| $A$ | – | 15 |
| $\mu$ | – | 50 |

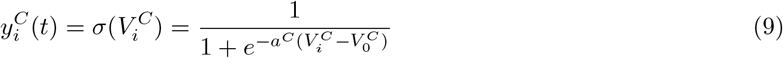

where *i* ∈ *{F, E}* The outputs of the cortical neurons are then subjected to a threshold check to implement WTA mechanism. For the cortical neurons, the parameters *a*^*C*^ = 8 and 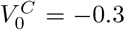 were used, while the threshold `thr` was set to 0.2.

#### Input signals to the motor cortex

The input signals to the neural oscillator are a key component of the model and were also used to analyze the system behavior in closed loop. These signals can be expressed as

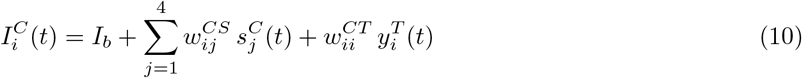

where *I*_*b*_ = 0.65 is a constant term used to sustain oscillations in the motor cortex oscillator. The coefficients 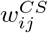 and 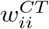 represent, respectively, the synaptic weights of the discrete sensory input signals (see Table 1) and the synaptic weight of the thalamus-to-cortex connection (see Table 5). Here, 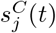 denotes the *j*-th element of the sensory input vector, and 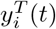 is the output of the *i*-th thalamic neuron.

#### GO part of the striatum

The Go neurons follow the dynamics described in [20], which is briefly recalled here and illustrated in Figure 2. Specifically:

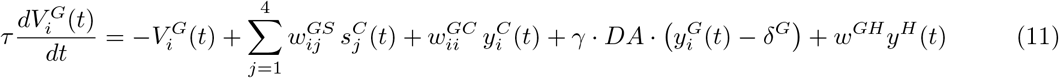

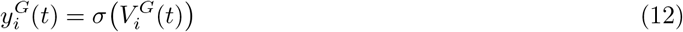

where 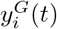 is the output of the *i*-th neuron in the Go part of the striatum. Each Go neuron receives excitatory input from the sensory cortex through the synaptic weights *W*_*GS*_ (see Table 1) and an additional excitatory projection from the corresponding cortical neuron via 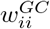 (see Table 5). The activity of Go neurons is modulated by the dopamine level *DA* (scaled by *γ >* 0) and by the activity of cholinergic interneurons *y*^*H*^ (*t*). Dopamine acts in a contrast-enhancing manner: it is excitatory if the neuronal output exceeds the threshold *δ*^*G*^ and inhibitory otherwise, thereby adding complexity to the WTA mechanism in the decision-making process. Conversely, the cholinergic interneurons are always inhibitory to the Go neurons given *w*^*GH*^ *<* 0 (see Table 5).

#### NoGo part of the striatum

In a similar manner, the dynamics of the NoGo neurons can be described as:

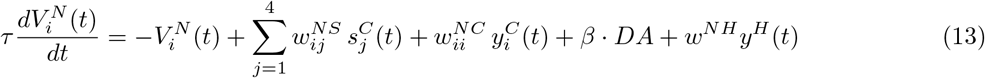

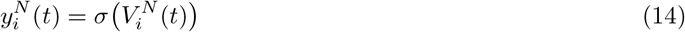

where 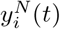 represents the output of the *i*-th neuron (Down or Up) in the NoGo compartment of the BG circuitry. According to the scheme in Figure 2, these neurons receive excitatory projections from the sensory cortex through the synaptic weights 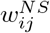 (see Table 1) and additional excitatory input from the corresponding cortical neurons via 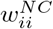 (see Table 5). As in the Go compartment, the dopamine level *DA* also modulates the dynamics of the NoGo neurons, but here it provides a constant inhibitory effect because *β <* 0. In contrast, the cholinergic interneurons provide excitatory input to the NoGo neurons since *w*^*NH*^ *>* 0 [20].

#### Globus pallidus pars externa

The equations governing the *F* and *E* neurons of this compartment are characterized by the following dynamics:

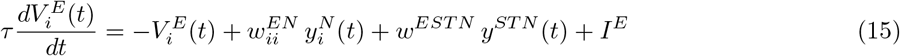

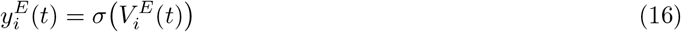

where 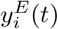 is the output of the *i*-th GPe neuron.Each neuron receives an inhibitory projection from the corresponding neuron in the NoGo compartment 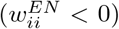 and an excitatory projection from the STN (*w*^*ESTN*^ *>* 0), as summarized in Table 5.

#### Globus pallidus pars interna

The equations governing the *D* and *U* neurons of this compartment follow the dynamics:

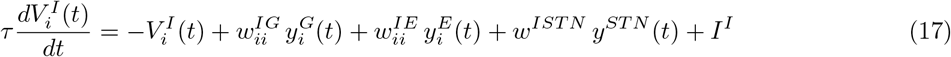

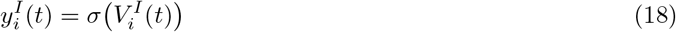

where 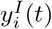 is the output of the *i*-th GPi neuron. Each neuron receives inhibitory projections from the corresponding neuron in the Go compartment 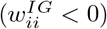 and from the 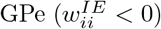. Conversely, excitatory input is provided by the STN compartment through the hyperdirect pathway (*w*^*IST N*^ *>* 0), as summarized in Table 5.

#### Subthalamic Nucleus

The activity of the STN was modeled using a single lumped dynamic representation, as reported in [20, 27]. The output of this compartment, *y*^*ST N*^ (*t*), and its corresponding state, *V* ^*ST N*^ (*t*), follow the equations:

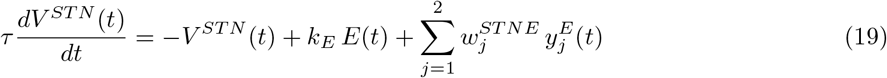

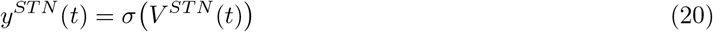

where the energy function is defined as

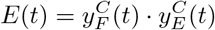

The STN is connected with the motor cortex and encodes the level of conflict in the action selection process through the energy function *E*(*t*). The strength of this conflict-monitoring mechanism is regulated by the parameter *k*_*E*_ (see Table 5). As illustrated in Figure 2, the STN also receives inhibitory projections from the GPe via the synapses 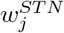.

#### Thalamus

The dynamics of the neurons in this compartment can be expressed as:

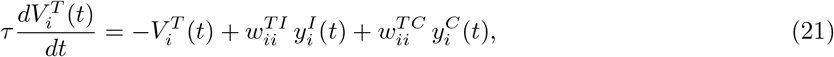

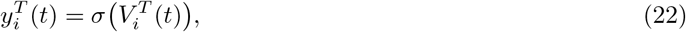

where 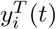 represents the output of the *i*-th thalamic neuron. Each neuron in this compartment receives an excitatory projection from the corresponding neuron of the motor cortex 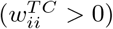 and an inhibitory projection from the corresponding neuron of the GPi 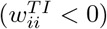. The synaptic parameters are reported in Table 5.

#### Cholinergic Interneurons

As for the STN, the cholinergic interneurons were modeled using a single differential equation. Let *y*^*H*^ (*t*) and *V* ^*H*^ (*t*) denote the output and state of this compartment, respectively. The dynamics can be expressed as:

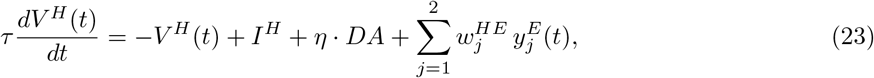

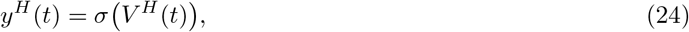

where the cholinergic interneuron compartment is inhibited by dopamine (*η <* 0) and remains active under basal conditions due to the constant input *I*^*H*^ . The full list of parameters is reported in Table 5.

### 5.8 Model Simulation Scheme

In order to perform model simulations, a forward Euler–Maruyama scheme was employed [15, 73]. The entire neuromechanical system was rewritten in the form of an autonomous stochastic differential system. By discretizing it, the following recursive scheme was obtained:

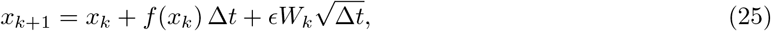

where *x*_*k*_ denotes the state vector at the *k*-th time step, *f* (*x*_*k*_) is the deterministic drift term, and Δ*t* is the integration step. The parameter *ϵ* defines the amplitude of the additive noise, while *W*_*k*_ are independent Gaussian random variables distributed as *N*(0, 1). for all the simulations *ϵ* was set equal 0.01, while the time step Δ*t* of 0.0001 s was adopted for discretization.

## Acknowledgments

Andrea Tigrini acknowledges the Japan Society for the Promotion of Science (JSPS) for the financial support received through fellowship grant PE25015.

